# Generating virtual patient data for in silico clinical trials of medical devices during extracorporeal membrane oxygenation treatment

**DOI:** 10.1101/2024.09.03.610823

**Authors:** Micha Landoll, Yifei Huang, Filippo Follegot, Stephan Strassmann, Ulrich Steinseifer, Christian Karagiannidis, Michael Neidlin

## Abstract

This study introduces a virtual patient generation model as online tool through the generation of high-quality synthetic data, addressing challenges like privacy cocerns and limited dataset sizes. Using a Conditional Tabular Generative Adversarial Network (CTGAN), we generated synthetic data from the Electronic Health Records (EHR) of 767 veno-venous extracorporeal membrane oxygenation (ECMO) patients, focusing on 55 critical therapy parameters. Rigorous preprocessing, imputation, and model tuning ensured that the synthetic data closely mirrored real patient records, achieving a 86.6% coverage score and minimal deviations in data correlations. The tool is integrated into a web platform - https://cve-sim.de/ecmo-vpg, allowing researchers to generate and visualize virtual patient cohorts, potentially enhancing ECMO research and reducing the need for extensive clinical trials.

## 1 Introduction

Clinical trials play a crucial role in the evaluation and approval of medium-to-high and high-risk medical devices [1]. However, the recruitment of eligible participants for clinical trials can pose challenges, particularly when seeking individuals with specific disease profiles, family histories, or lifestyles that align with the trial’s inclusion criteria [2]. The increasing complexity of regulations, financial demands, and ethical considerations further contribute to the challenges faced by traditional clinical trials [3].

In this regard, there has been an interest to use existing clinical data for analysis of disease phenotypes and evaluation of treatments [4]. Specifically, electronic health records (EHRs) have become an increasingly common approach for studying real-world patient data. However EHRs are limited by their shortcomings inherent to its complexity and retrospective nature. For instance, time-stamped observational data from EHRs may appear as unrealistic during retrospective analysis and certain inclusion criteria for patients may introduce sample selection biases which can result in misleading results [5].

In light of these challenges, the question arises: Is it possible to generate synthetic data from EHR databases to directly contribute to clinical evidence?

These virtual patients can then be integrated in computer simulations, facilitating the use of in-silico clinical trials (ISCTs) for the development or regulatory evaluation of medical devices or clinical interventions [6]. This approach has the potential to significantly reduce the risk to patients, as well as the time and cost associated with traditional clinical studies. Furthermore, a virtual patient cohort could supplement traditional clinical trials by decreasing the required number of enrolled patients, enhancing statistical significance, and providing a safer alternative. Such virtual patient data could also offer insights to inform clinical decision-making [7]. Additionally, the virtual patient cohort can circumvent the need to access sensitive personal information stored in EHR data [8]. One application that would benefit from further patient-specific insights is extracorporeal membrane oxygenation (ECMO). The intensive treatment offers functional support for respiratory and hemodynamic functions through the use of an artificial heart pump and an artificial membrane lung. It has emerged as a promising intervention for patients experiencing severe cardio-respiratory failure that do not respond to conventional treatments [9]. However, its significant technical requirements and associated risks continue to restrict its widespread application [10, 11]. The nuanced understanding of patient-device interaction during ECMO therapy is a major challenge to enable exploration of various design and therapy settings for ECMO device developers and hence bring novel medical devices to market. Many computational models of ECMO exist in the literature, however those are always run on “characteristic patients” or fitted to few datapoints. For example, the hemodynamics of the cardiovascular system during ECMO have been represented in-silico by lumped parameter modeling [12, 13, 14] and validated by hydraulic in-vitro experiments [13]. Virtual patient cohorts providing realistic conditions for the computational models would thus push their capabilities in terms of application and prediction quality.

There are two main methodological categories for generating synthetic data. The first involves generating numbers from distributions using statistical methods, while the second employs agent-based or artificial intelligence based modeling methods based on deep learning models, like variational autoencoders (VAEs) and generative adversarial networks (GANs) [4]. The idea of generating synthetic data for virtual patients has been around since its first description published in 1971 [15, 16] and there exist numerous examples. Korchani et al. developed a tool generate synthetic data from summary statistics of real-world clinical data and provide users with the capability to interactively optimize the synthetic dataset [4]. Ganguli et al. developed a novel generative recurrent neural network framework to produce accurate, applicable, and de-identified synthetic medical data for patients with metastatic cancer [17]. Romero and Lozano showed that GANs can produce reliable, clinically-driven cohorts of thoracic aortas with good efficiency and proposed the integration of machine learning models to expedite the process of cohort generation [18]. In general, GANs have demonstrated to be successful in producing complex synthetic data, but they often suffer from mode collapse, limited diversity in generated synthetic datasets, and inaccurately generated data [17].

The evaluation of synthetic data quality is a major challenge in data generation tasks [19]. A sufficient overlap with the underlying real-world data has to be shown, which can be challenging with high-dimensional and multi-variate datasets. In addition, further downstream integration into computational models should produce indistinguishable predictions whether the models are run on synthetic or real-world data. This depends very much on the specific question that is investigated with the model and needs to be checked on an individual basis. Thus any virtual patient generator should provide an easy way to be shareable and directly usable by the scientific community, especially researchers working in computational modelling of ECMO.

The aim of this study is to develop a virtual patient generation tool based on real-world data from ECMO patients using a conditional tabular generative adversarial network (CTGAN). Data quality will be assessed and the integration of the tool in a web-based platform will be presented.

## 2 Materials and Methods

### 2.1 Patient data

Data from EHRs of 598 ARDS patients who underwent veno-venous ECMO treatment within the time frame between Oktober 2012 and September 2023 was used. The study has been approved by the Institutional Review Boards of the University Hospital Aachen and the ARDS- and ECMO Centre in Cologne-Merheim (Witten/Herdecke University). A summary of the cohort is shown in Table 1

**Table 1:** Description of the patient cohort regarding demographics and treatment.

|  | Count | Mean | SD | Min. | 25<sup>th</sup> | Median | 75<sup>th</sup> | Max. |
| --- | --- | --- | --- | --- | --- | --- | --- | --- |
| Age (years) | 598 (100%) | 54.5 | 12.9 | 18.3 | 46.5 | 57 | 64 | 85.9 |
| BMI | 331 (55.3%) | 31.2 | 10.1 | 13 | 25.2 | 28.7 | 34.7 | 97.3 |
| Female proportion | 196 (32.8%) | - | - | - | - | - | - | - |
| Male proportion | 402 (67.2%) | - | - | - | - | - | - | - |
| ICU length of stay (days) | 598 (100%) | 31.8 | 26.9 | 1 | 15 | 24 | 42 | 210 |
| ECMO duration (hours) | 598 (100%) | 561.2 | 564.5 | 0 | 229.3 | 401.5 | 690 | 4973 |
| Survival | 337 (56.4%) | - | - | - | - | - | - | - |

Two datasets were extracted for further analysis.

- Low-dimensional dataset A of patients (n=76) who underwent Pulse Contour Cardiac Output (PiCCO) monitoring during ECMO.
- High-dimensional dataset B of all patients (n=767) who underwent ECMO treatment

In both cases, features with more that 21% missing values were removed and all time points were considered at once. Each patient was described by a unique case ID. This yielded 8 features and 35,293 data points in the low-dimensional high temporal resolution dataset A (Table 2) and 55 features and 14,997 data points in the high-dimensional low temporal resolution on daily base in dataset B. Missing data was imputed using the hyperimpute package, which is an iterative imputer using both regression and classification methods based on linear models, trees, XGBoost, CatBoost, and neural nets [20]. The chosen imputation for the PICCO datasset A settings are presented in supplementary Table S1. The high dimensional dataset B has been imputed using a multivariate imputation by chained equations (MICE) using the *sklearn*.*IterativeImputer* function. The histograms of all features after data pre-processing can be found in Figures S1 and S2.

**Table 2:** List of features of the low-dimensional dataset A.

| Parameter | Parameter | Parameter | Parameter |
| --- | --- | --- | --- |
| HR | IBPm | PAdia | PAsys |
| IBPdia | IBPsys | PAm | Respiratory rate |

### 2.2 Synthetic data generation

Conditional Tabular Generative Adversarial Networks (CTGANs) were used for the generation of synthetic data [21]. The basic idea behind GAN is to simultaneously train two models: a generator and a discriminator. The generator, which is commonly depicted as a deep neural network, creates new data instances from random noise as input that resemble the training data. The generator continuously adjusts its output to produce samples that closely mimic real data as being trained using back propagation to tune its parameters. The discriminator functions as a binary classifier which is responsible for differentiating between generated and original input. These two models contest with each other and iteratively improve their respective performance through an adversarial training process, resulting in the generation of increasingly realistic synthetic data. [22].

The data set was split into 80 % training and 20 % test data, stratified by patient level to avoid data leakage. Data leakage may cause invalid machine learning models due to over-optimization of the model and it arises because data or information beyond the training set is incorporated during the learning process [23]. In the context of the clinical data derived from the EHRs collected at different time intervals from individual patients, there exists a time-dependent internal relationship between data points.

The CTGAN from the *synthcity* package was used including hyperparameter optimization with the Optuna package. The AUCROC curve of the discriminator was minimized in the training process. This resultred in a network with the hyperparameters as shown in Table 3.

**Table 3:** Best set hyperparameters of the CTGAN model.

| Hyperparameter | Value |
| --- | --- |
| generator_n_layers_hidden | 4 |
| generator_n_units_hidden | 100 |
| generator_nonlin | 'leaky_relu' |
| n_iter | 1000 |
| generator_dropout | 0.1159 |
| discriminator_n_layers_hidden | 1 |
| discriminator_n_units_hidden | 100 |
| discriminator_nonlin | 'elu' |
| discriminator_n_iter | 5 |
| discriminator_dropout | 0.0503 |
| lr (learning rate) | 0.0002 |
| weight_decay | 0.001 |
| batch_size | 100 |
| encoder_max_clusters | 9 |

### 2.3 Evaluation

As the Generator model of the CTGAN is a neural network, it is considered a black box model in interpretabitility. Therefore an indirect evaluation must be considered. The quality of the synthetic data was evaluated in several ways. At first, the following three scores, ranging from worst (0) to best (0), were used for each feature to assess the quality. Each property ranges describes:

- Coverage - whether a synthetic column covers the full range of values that are present in a real column
- Boundary - whether a synthetic column respects the minimum and maximum values of the real column
- Synthesis - whether each row in the synthetic data is novel, or whether it exactly matches an original row in the real data

Additionally, bivariate and multivariate analyses were performed. In the bivariate case, Pearson’s pairwise correlation and absolute differences in the correlation matrices between synthetic and real data were compared. In the multivariate case, the t-distributed Stochastic Neighbor Embedding (t-SNE), Uniform Manifold Approximation and Projection for Dimension Reduction (UMAP) and Principal Component Analysis (PCA) was employed. T-SNE is a technique to visualize high-dimensional data in two-dimension using non-linear dimensionality reduction.

### 2.4 Web-based visualization tool

The trained model was implemented in a web-based graphic user interface using the online dash tool [24]. It serves as a stratification, generation and visualization tool of the virtual patient cohort with its main functions listed as follows:

- Cohort selection based on demographic features
- User input text box for sample number to be generated
- Integrated CTGAN model generate data samples
- Data is filtered by selected demographic cohort characteristic
- Visualized feature distribution, correlations and comparisons between original and synthetic data
- Export of synthetic data to CSV file

## 3 Results

Coverage scores of all features for dataset A are shown in Figure 1 a) in an ascending order, ranging from 0.76 to 0.97. With an overall average coverage score of 0.90, the synthetic dataset proves to be capable of covering approximately 90% of the numerical ranges in the original dataset in average. The other two scores, representing boundary and synthesis, are both 1.0. Thus, the synthetic data adheres to the minimal and maximal boundaries of the original data. The synthesis score 1.0 means that all of the synthetic data points are not merely pure copies of the original data points but rather novel ones generated by the CTGAN model. The same analysis was expanded to the high-dimensional dataset B, see Figure 1 b). The synthetic dataset’s coverage of the original dataset ranges from 56% to 100% with an overall average of 86.6%. Synthesis and boundary scores remain at 1.0.

**Figure 1:**
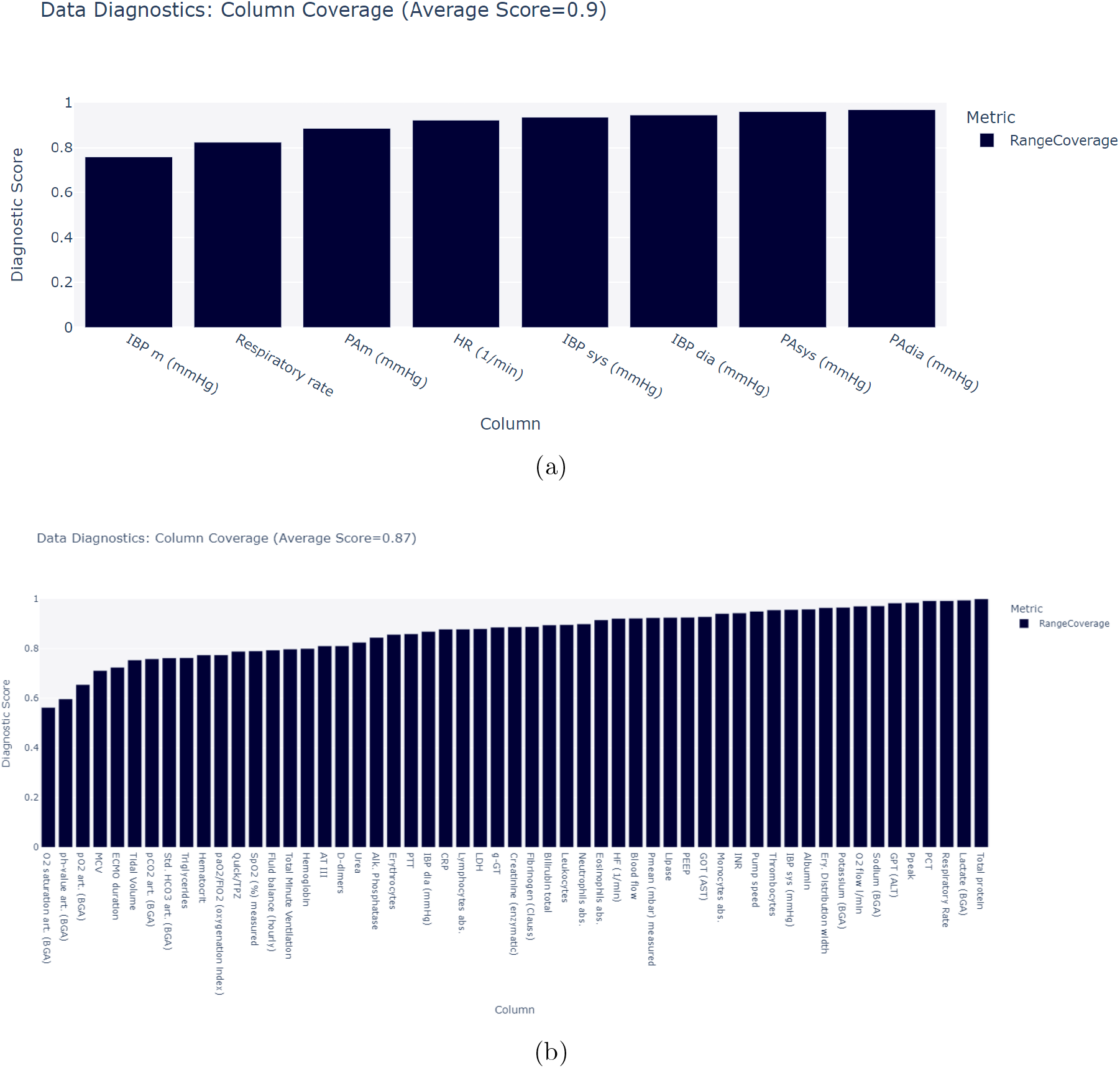
Coverage scores for (a) dataset A and (b) dataset B

Further comparison through histograms is presented in Figure 2 for each feature. The synthetic data maintains a close alignment within the boundaries of each feature in the original data as already described by the boundary score of 1.0. However, the area under the curve reveals that the synthetic data does not cover the full range of the original data. This corresponds with the coverage score of 0.90, signifying that the synthetic dataset covers approximately 90% of the numerical ranges in the original dataset on average.

**Figure 2:**
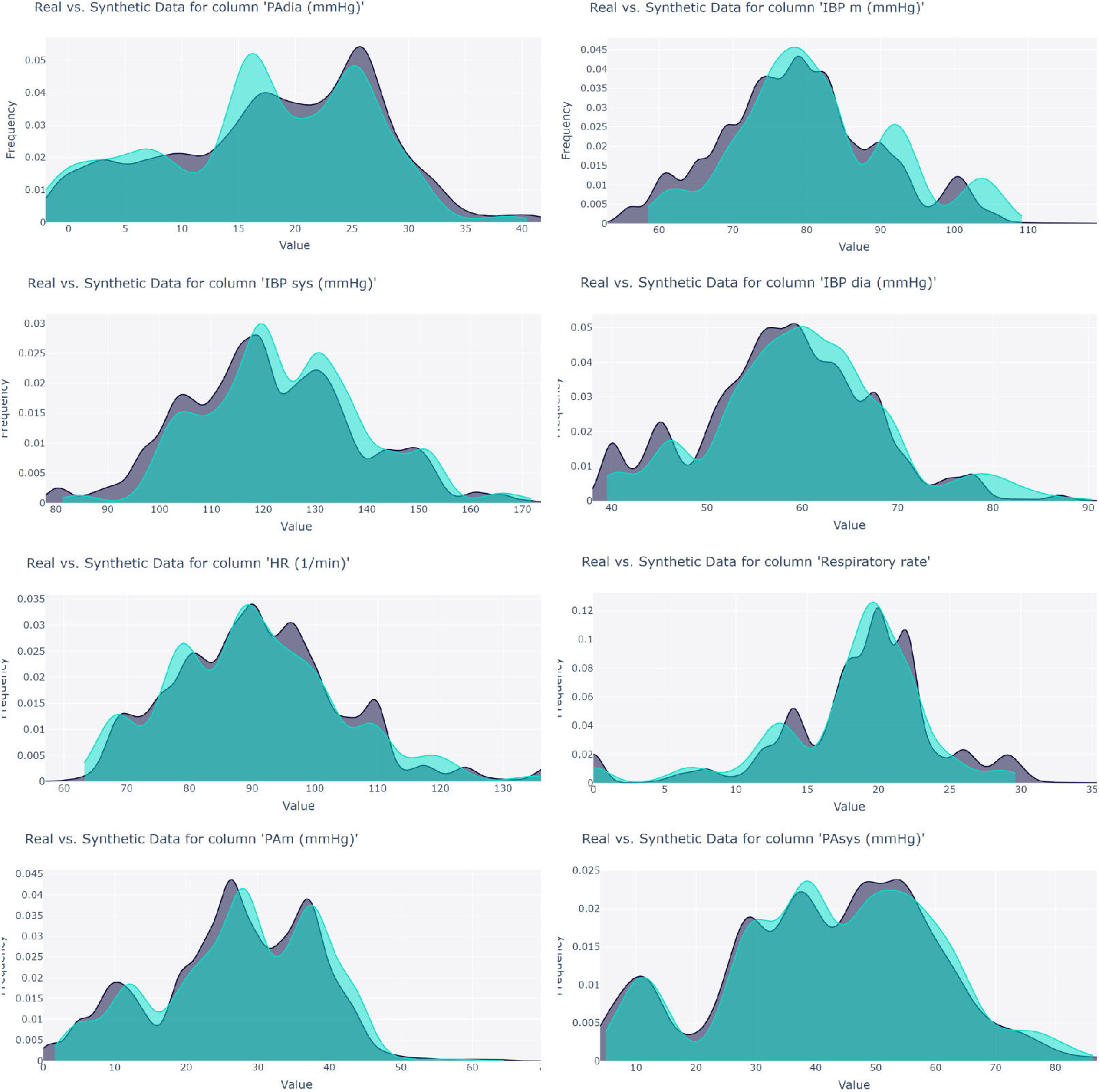
Curve plots illustrating the distribution of values for each feature in the synthetic dataset (cyan curves) against the original dataset (dark grey curves)

### Bivariate comparison

In a pairwise comparison we calculated the monotonic correlation. As metrics we used correlation by Pearson for dataset A and correlation by Spearman for dataset B. The correlation matrices for dataset A of both original and synthetic data are shown in Figure 3. The correlation values are calculated and displayed in an absolute range of [0, 1]. The color map of both Pearson matrices offers visual insights of the degree of similarities between synthetic and original data feature-wise. Overall, the pairwise relationships between the features are preserved in the synthetic dataset. For a more quantitative comparison, the mean values of absolute correlation difference for each feature range from 0.04-0.08, with an overall average mean of 0.059. The maximal absolute difference is 0.16, which appears between the feature pair *PAm (mean pulmonary artery pressure)* and *PAdia (diastolic pulmonary artery pressure)*. The absolute Spearmans correlation for dataset B for each feature combination is shown for the real dataset in Figure S3 and the generated cohort in Figure S4. The absolute difference in Spearman’s correlation is shown in Figure S5 and highlights significant differences of some features (GOT, GPT, LDH), with a maximum discrepancy of 0.35. Nevertheless, calculating the mean of the absolute correlation differences on a feature-wise basis, yields relatively low differences. Furthermore, averaging the means of absolute correlation differences across all features yields a discrepancy of *≈* 0.032. Furthermore the mean correlation difference for each feature is presented in Figure 4. Most of the monotonic intervariable relationships are preserved in the synthetic data, but some combined treatment effects may be underrepresented.

**Figure 3:**
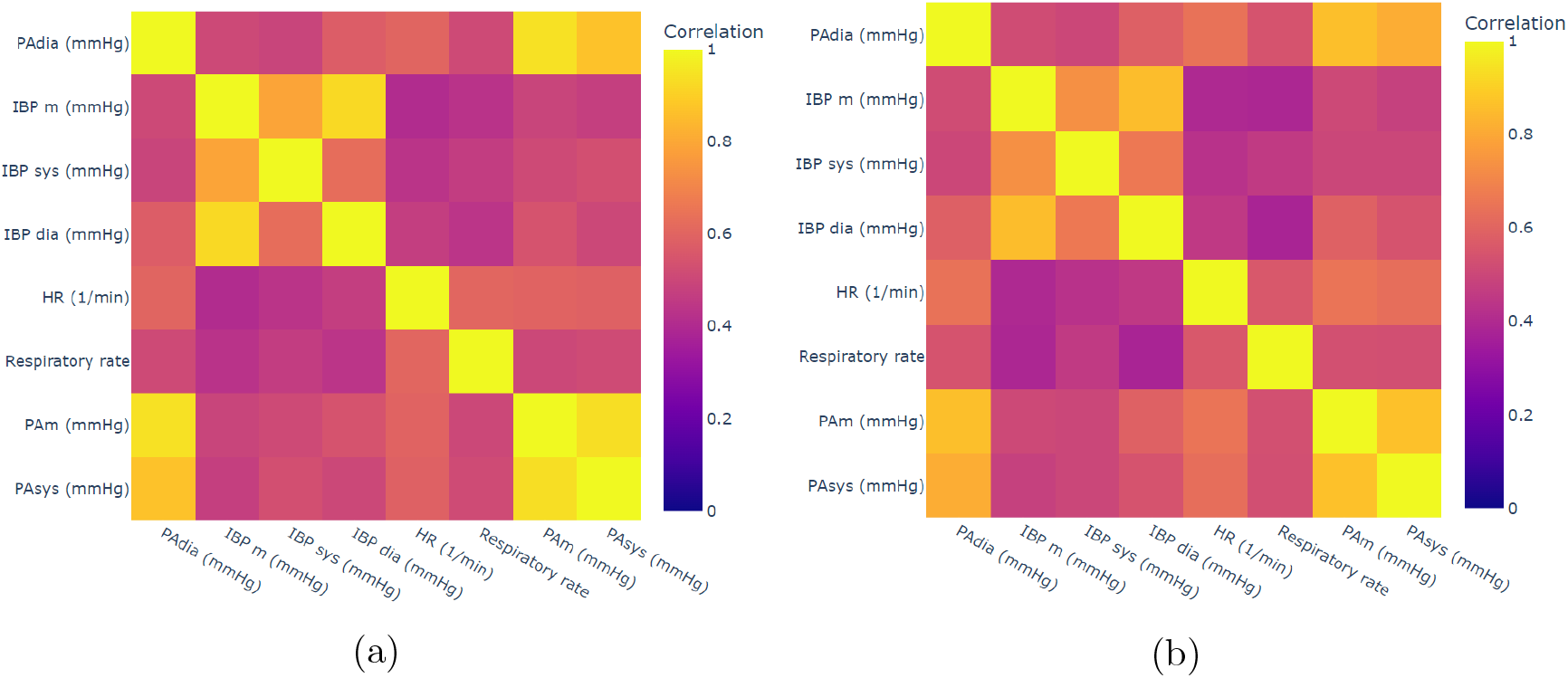
Absolute Pearson correlation matrix of (a) original and (b) synthetic PICCO datasets

**Figure 4:**
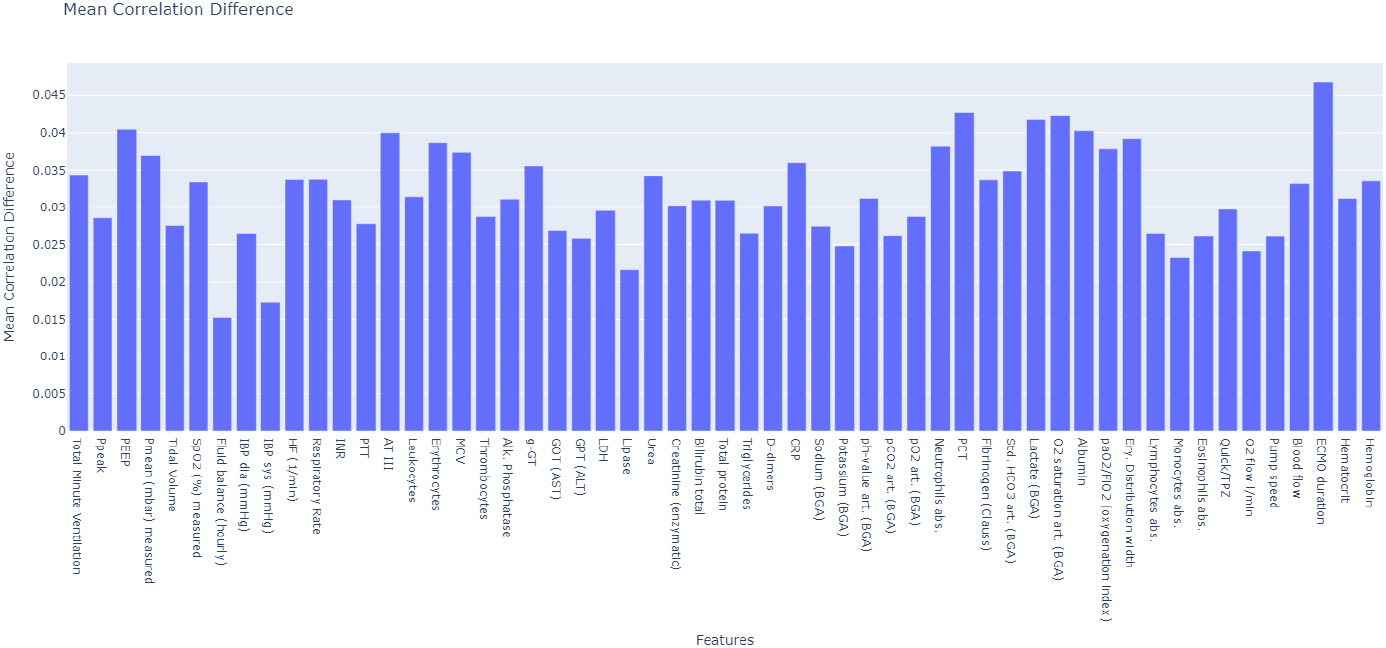
Feature-wise mean correlation difference between real and synthetic data.

### Multivariate comparison

Figure 5 a) illustrates the t-SNE plot of both original and synthetic data for dataset A. It is important to note that the distances between points in this low-dimensional plane represent similarities rather than directly comparable distances. It can be observed from the t-SNE plot and its dimensional distributions that the synthetic data points in orange are encapsulated within the scope of original data points in blue, with a significantly higher density of synthetic data points in the central region of the plot.

**Figure 5:**
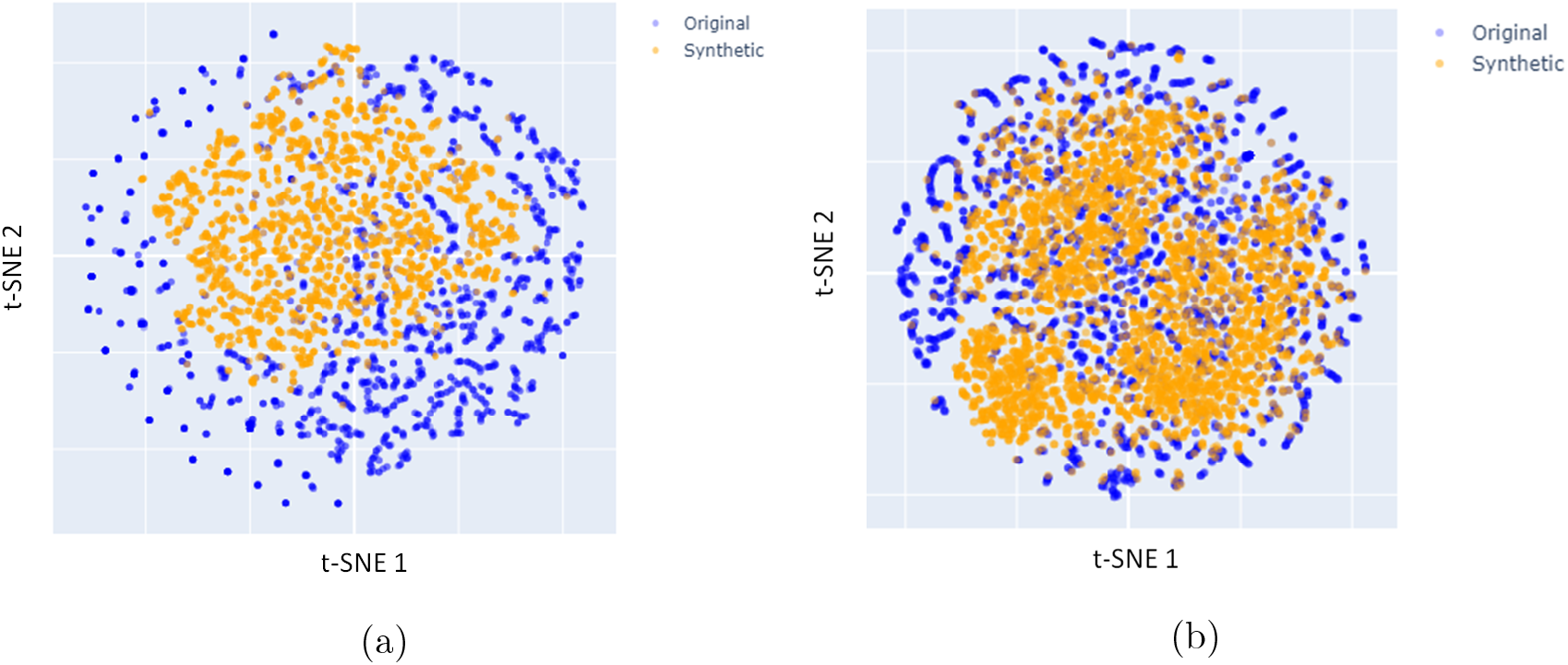
t-SNE of original (blue) and synthetic (orange) data with dimensional distributions of both axes for dataset A (a) and dataset B (b)

The t-SNE plot of both original and synthetic data of the-dimensional dataset B is presented in Figure 5 b). Similar to the one of the low-dimensional dataset, synthetic data points in orange are included within the range of original data points in blue, strictly complying with the boundaries set by the original data points.

### 3.1 Visualization tool

This section gives insights into the functionality and layout of the web visualization tool developed using *Dash*, as illustrated in Figure 6. The dashboard is divided into two sections: cohort selection and synthetic data generation.

**Figure 6:**
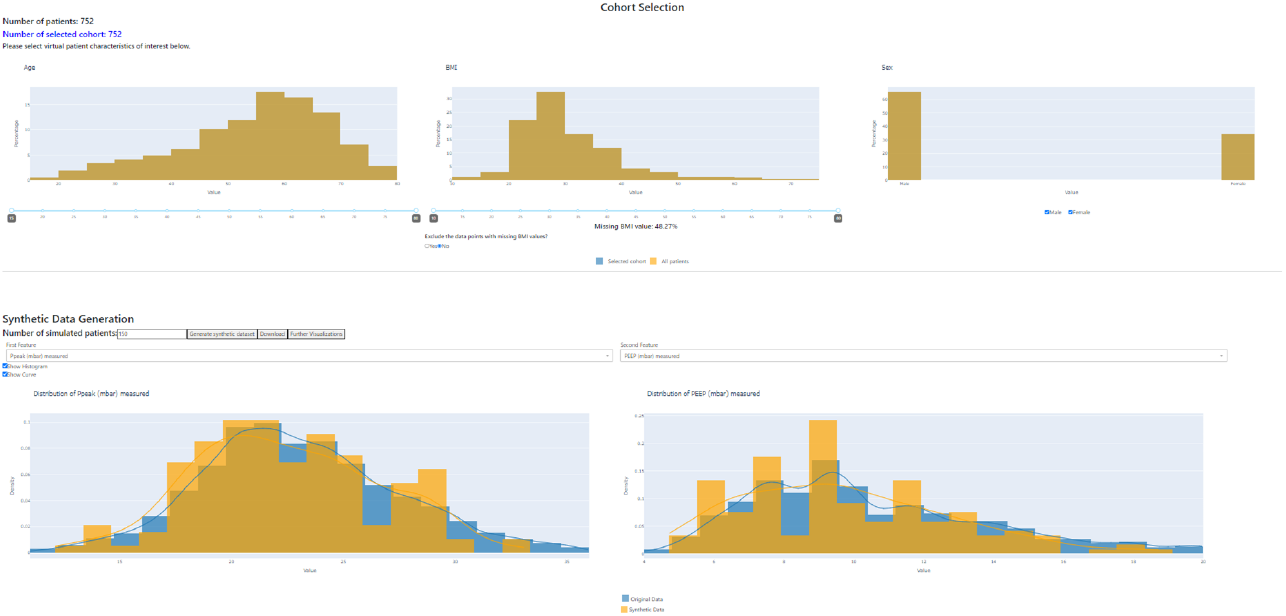
A sample overview of the visualization tool in Dash

The top section of the dashboard is dedicated to cohort selection on demographic parameters, offering users an overview of the patient cohort from the EHR database and selection of subcohorts based on BMI, age, and gender.

The lower section of the visualization tool is dedicated to the generation and visualization of synthetic data. In the generation process the datapoints are assessed if they match the distributions of the selected cohort characteristics. Neglected datapoints will be iteratively resampled until the desired size of the generated cohort is met. The generated synthetic data is then visualized and compared through histograms and simplified distribution curves of each feature. Blue represents stratified original data, while orange represents synthetic data. At the end of this section, the user can download the generated virtual patient cohort as a CSV file. Each column in the file corresponds to a specific feature. The virtual patient generator can be free accessed through https://cve-sim.de/ecmo-vpg.

## 4 Discussion

In this work, a virtual patient generation tool has been developed to synthesize data from the EHR database of 598 V-V ECMO patients utilizing a Conditional Tabular Generative Adversarial Network (CTGAN). The CTGAN model successfully captured the statistical distributions of the original EHR data, generating synthetic datasets with similar structure and inter-variable relationships. Validation of the accuracy and quality of the synthetic data was performed using various metrics, including uni-, bi- and multivariate comparisons. In both, a high-temporal resolution and low-dimensional dataset (dataset A) and a low-temporal resultion and high-dimensional dataset (dataset B) we were able to create datapoints that effectively mirror the structure and internal relationships in the original data. We have observed that the model can reflect the statistical characteristics of the original dataset. Furthermore we saw, that the synthetic data is more than just a copy of training data and may be a valuable contribution to solve privacy issues of patient data. While the univariate and bivariate analyses have shown a very good overlap between real and synthetic data, the t-SNE embedding has uncovered that there are regions not described by the synthetic data. Also in bivariate analysis we have discovered regions, where the combination of multiple complications and treatments effect may be underrepresentated in the synthetic data. More specifically, the synthetic data generator stayed within “conservative” bounds of the real data and did not generate datapoints close to the boundaries, see Figure 5. As a consequence, more uncommon patient situations might not be captured by the data generator. This observation underlines the need of a thorough understanding of the underlying patient data prior synthetic data generation. For instance identification of sub-cohorts (phenotypes) from the real-world data might support the choice of the ground truth for the GANs or the creation of individual models for the sub-cohorts.

Another critical observation was that obvious mistakes in the clinical data (some patients had larger diastolic than systolic pressures) can be also captured by the GANs. Hence, a careful analysis and data cleaning is required. In this study we did not consider the transient nature of the EHRs and all datapoints were put together. Time-series GANs [25] might offer a solution approach to consider the temporal dimension of the data.

To enhance user experience, the implemented CTGAN model has been seamlessly integrated with data querying, visualization of features, dynamic adjustment of settings and data exportation functionalities into a dashboard. The existing framework is able to provide data distributions for many parameters that are relevant for medical devices such as blood flows, pressures, blood gases and mechanical ventilator settings. Based on this data, virtual cohorts can be quickly created and implemented in computational models.

## A Appendix

### Settings of hyperimpute algorithm

**Table S1:** Keyword arguments of Hyperimpute.

| Keyword argument | Definition | Type | Parameterization |
| --- | --- | --- | --- |
| optimizer | The optimizer to use: "simple", "hyperband", "bayesian" | string | "hyperband" |
| classifier_seed | Model search pool for categorical columns | list | ["logistic_regression", "random_forest", "xgboost", "catboost"] |
| regression_seed | Model search pool for continuous columns | list | ["linear_regression", "random_forest_regressor", "xgboost_regressor", "catboost_regressor"] |
| class_threshold | The maximum number of unique items required for a column to be classified as a categorical column | integer | 5 |
| imputation_order | 0: ascending<br>1: descending<br>2: random | integer | 2 |
| n_inner_iter | Number of imputation iterations | integer | 10 |
| select_model_by_column | True: select a different model for each column;<br>False: reuse the model chosen for the first column | boolean | True |
| select_model_by_iteration | True: select new models for each iteration;<br>False: reuse the models chosen in the first iteration | boolean | True |
| select_lazy | True: if there is a trend in the current iteration, use the same model class for all columns without restarting the optimizer;<br>False: initiate the optimizer for each column unless other restrictions apply | boolean | True |
| select_patience | Number of iterations without objective function improvement to wait | integer | 5 |

#### Histograms of features

**Fig. S1:**
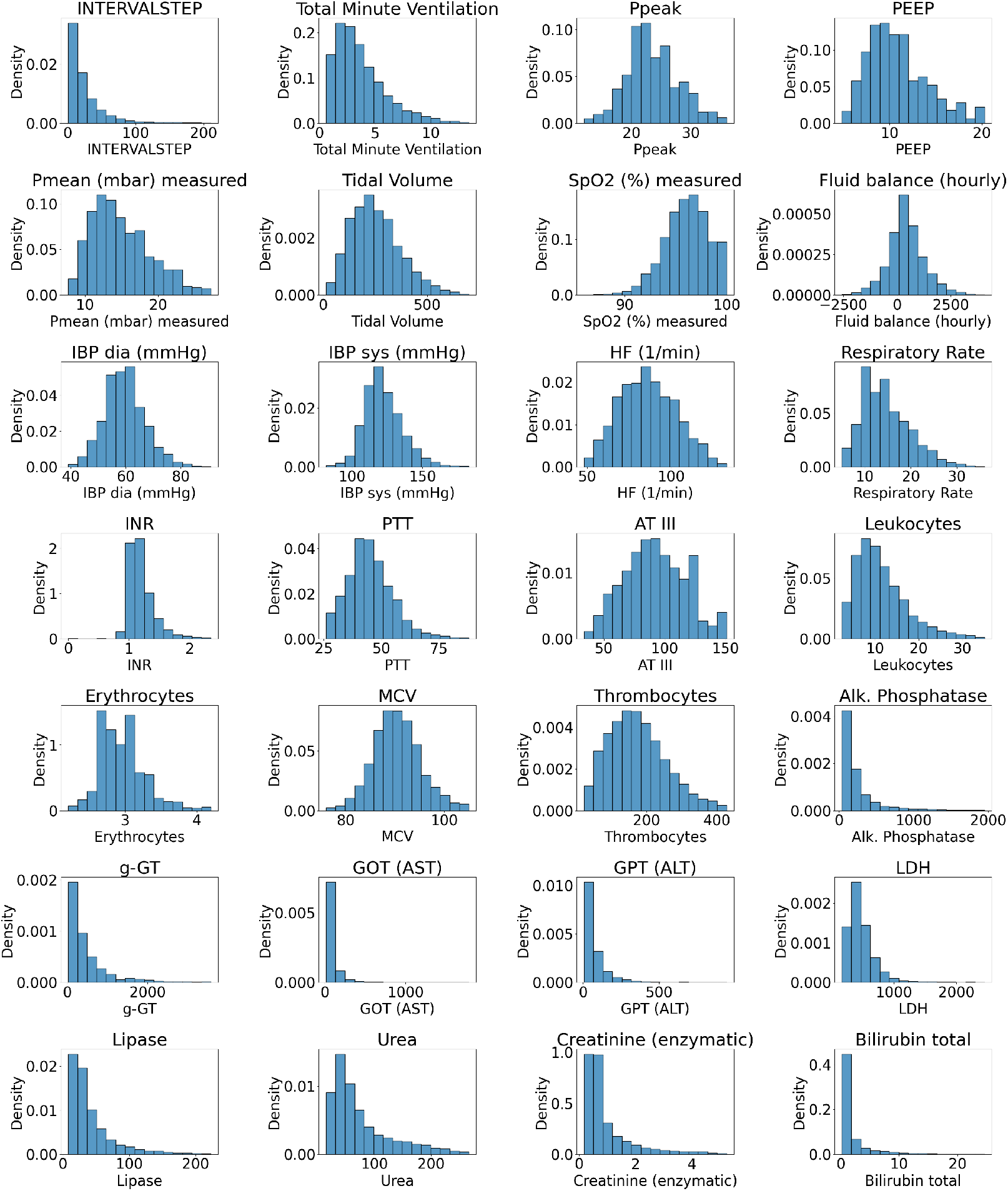
Histograms of all features (Part 1 of 2)

**Fig. S2:**
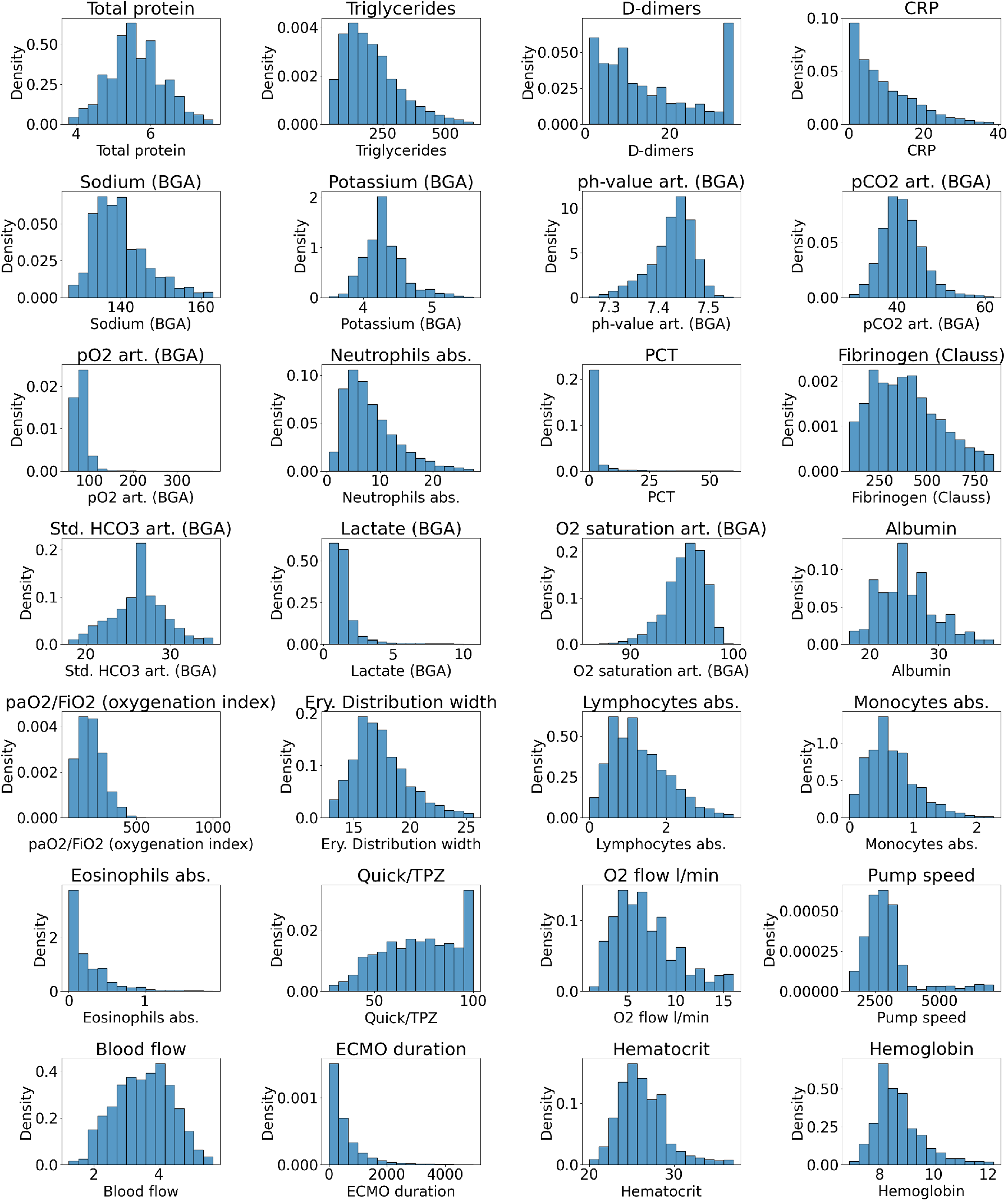
Histograms of all features (Part 2 of 2)

**Fig. S3:**
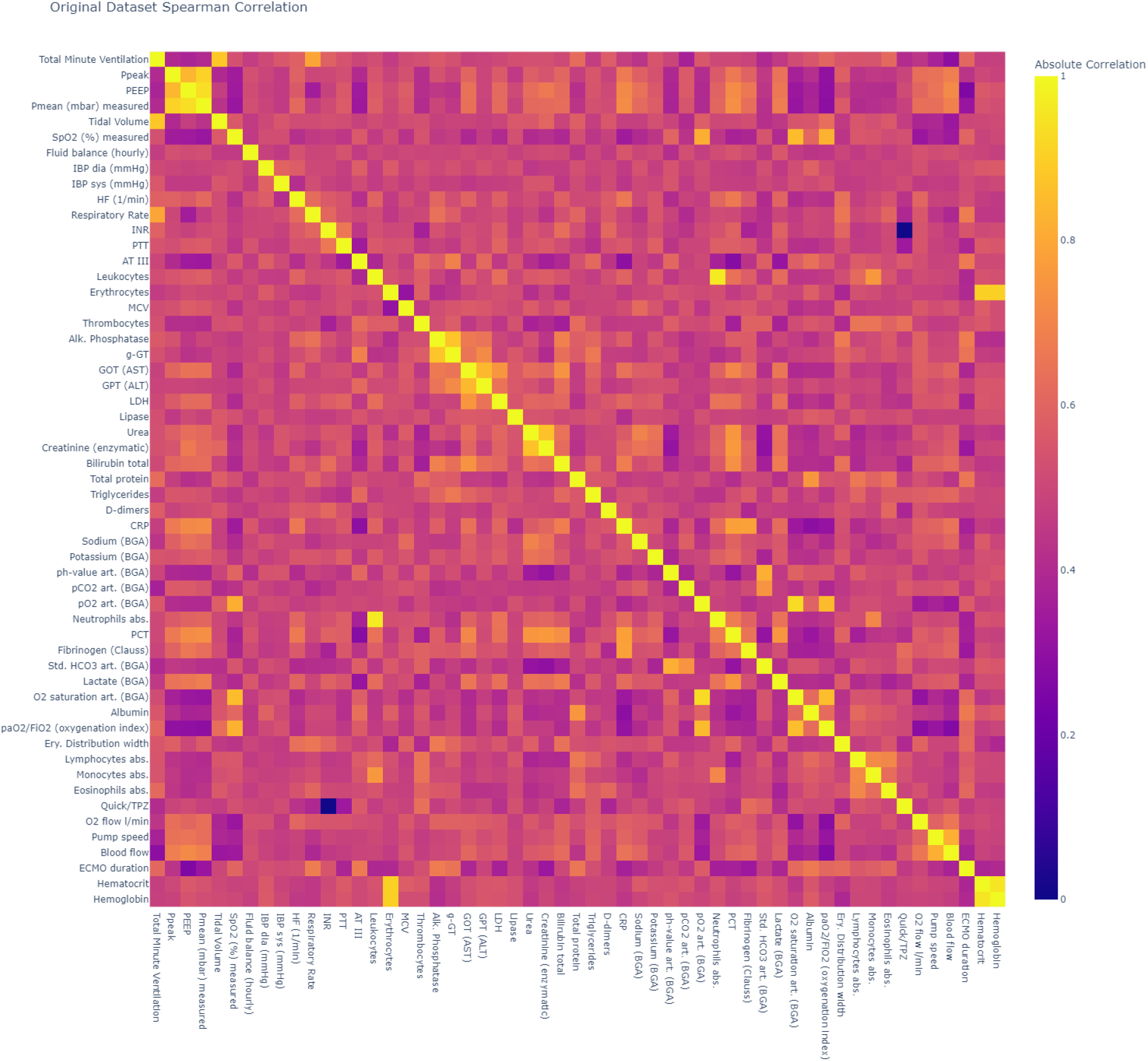
Absolute Spearman correlation of the original dataset B

**Fig. S4:**
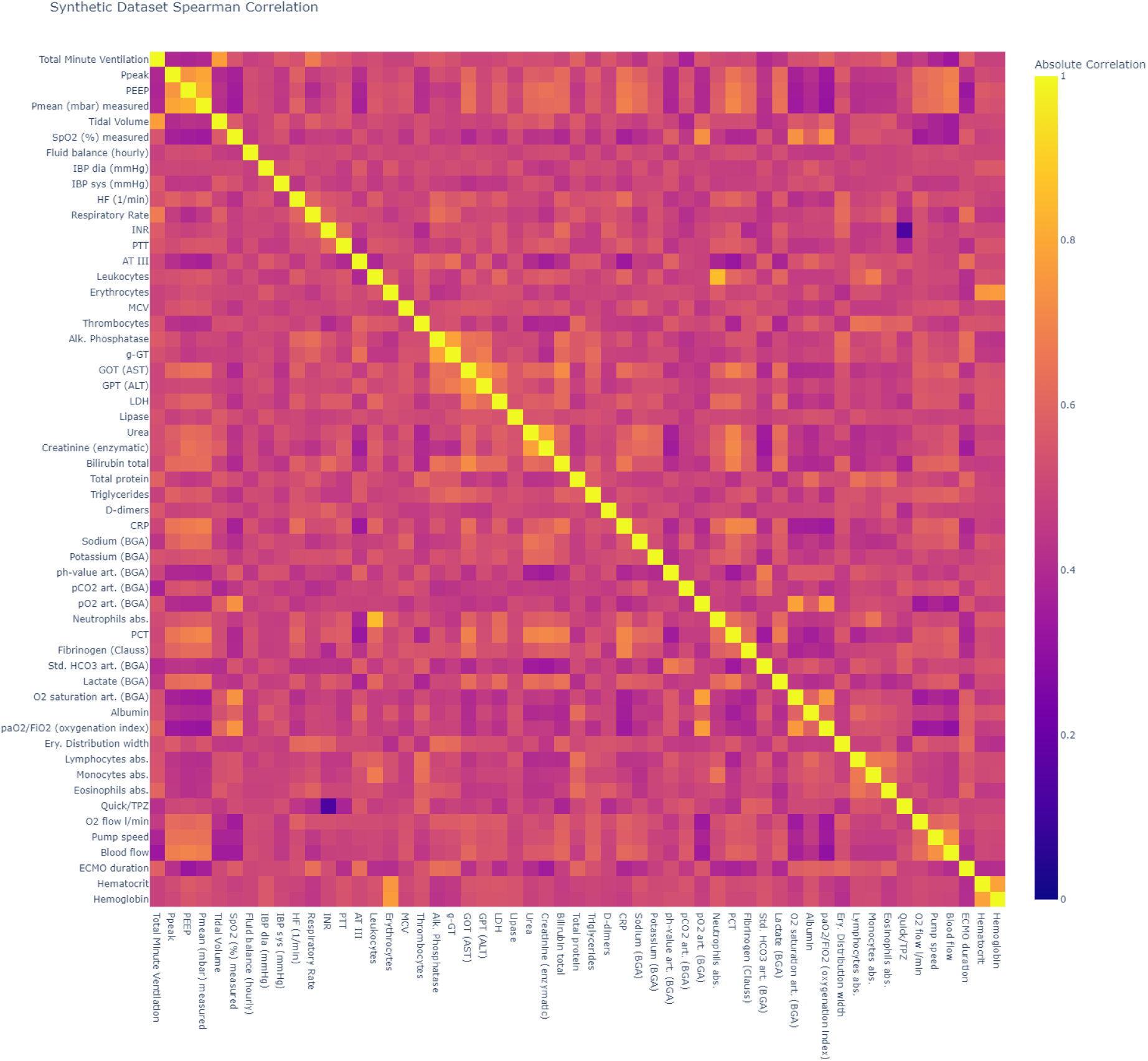
Absolute Spearman correlation of the synthetic dataset B

**Fig. S5:**
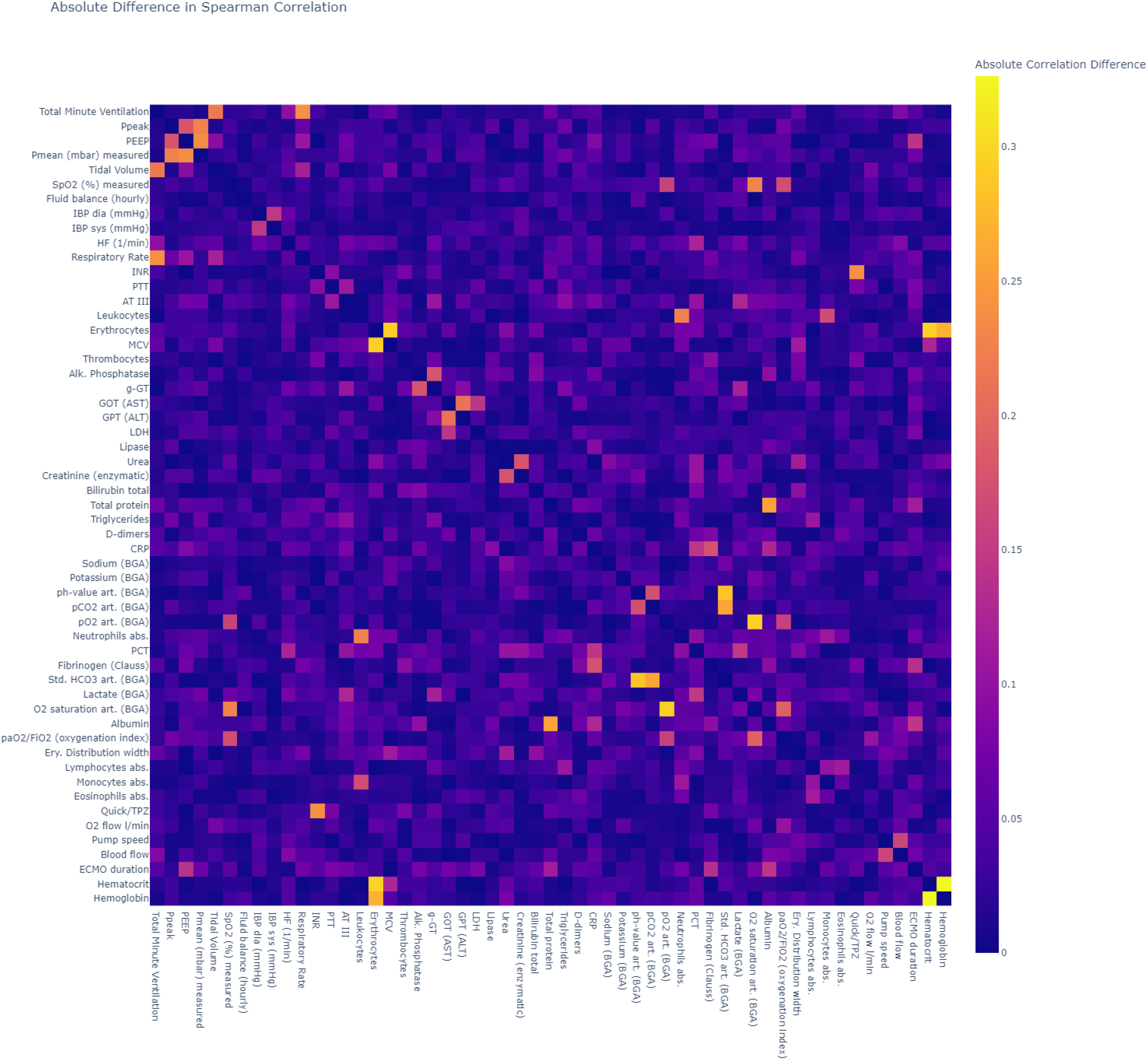
Absolute difference in Spearman correlation of the high-dimensional dataset B

